# Fair-weather friends: Unequal partnerships between *Parastagonospora nodorum* and *Pyrenophora tritici-repentis* define disease dynamics in wheat

**DOI:** 10.1101/2025.10.09.681059

**Authors:** Leon Lenzo, Evan John, Jason Bradley, Geoff Thomas, Dion Bennett, Kar-Chun Tan

## Abstract

*Parastagonospora nodorum* and *Pyrenophora tritici-repentis* are the causal agents of septoria nodorum blotch and tan spot of wheat, respectively. Though these fungal phytopathogens have been found to frequently cohabitate the same leaf, their interaction dynamics in the manifestation of disease remain poorly understood due to limitations in species-specific detection methods. We developed a digital PCR based model targeting conserved regions of the α-tubulin gene, enabling biomass quantification of both pathogens during infection. Field surveys revealed up to two in three symptomatic infections involved both pathogens, with co-infected plants showing significantly higher individual pathogen biomass than single-species infections. Host plants in the field with moderate resistance to both pathogens were found to be significantly more necrotic under co-infection, with individual pathogen biomass up to twice that observed value for single infections. However, like fair-weather friends the partnership between these two pathogens seems to be conditional.When *P. tritici-repentis*established first, secondary *P. nodorum*colonisationled to a breakdown of host resistance. Conversely, when *P. nodorum* established on the host first, it suppressed *P. tritici-repentis* colonisation regardless of host resistance. To our knowledge this is the first description of asymmetric priority effects overcoming host resistance in a plant pathosystem. Resistance breeding strategies evaluating single pathogen challenges may inadvertently select for cultivars vulnerable to sequential co-infection, necessitating integrated disease complex approaches for durable resistance development.

## Introduction

The order in which microorganisms colonise habitats fundamentally shapes community assembly and species interactions, a phenomenon known as priority effects ^1^. In the colonisation of plant tissues, early arriving species modify environmental conditions and resource availability, altering biotic conditions to determine the trajectory of community development ^2^. Agricultural phyllospheres represent a synthetic, simplifiedecological context, where vast expanses ofgenetically homogeneoushosts grown athighdensity create conditions that accelerate pathogen evolution andpromote rapid populationgrowth ^3^. Previouswork has demonstrated priority effects in plant-associated microbial communities, with pathogen arrival order determines infection outcomes and host responses, yet their operation in agricultural pathosystems remains poorly understood despite their potential to fundamentally alter disease outcomes ^4,5^. Recent surveys using molecular analysis across European wheat fields revealed that over two thirds of symptomatic leaf samples harboured at least two pathogenic fungal species ^6^. Environmental monitoring studies using spore trapping showed propagules from fungal pathogens circulate simultaneously during growing seasons, creating opportunities for co-infection ^7^. While priority effects have been extensively documented in bacterial pathosystems, with pathogens employing diverse strategies including competition, cooperation, and temporal separation to succeed in polymicrobial environments, their operation in fungal plant pathosystems remains poorly understood despite their potential to fundamentally alter disease outcomes ^8^.Polymicrobialinfections inplant systemsmayresult inresource competition, host immune modulation or direct pathogen-pathogen interactions that chemically or mechanically modulate the colonization, reproduction, and transmission success of coinfecting species ^9^. These interactions can be either facilitative, where one pathogen creates conditions that enhance proliferation of another ^10,11^, or inhibitory, where pathogens actively suppress competitors through antimicrobial compounds or resource monopolization ^12,13^.

*Parastagonospora nodorum*and *Pyrenophora tritici-repentis*, the causal agents of septoria nodorum blotch (SNB) and tan spot (TS) of wheat, represent an ideal system for investigating priority effects innecrotrophic plant pathosystems ^14,15^, with previous work documenting near ubiquitous co-occurrence across wheat growingregions ^16^.Empiricalstudiesof interactionsbetweenspecieshave been limitedhowever, largely due to the morphological similarity of their constituent diseases and lack of robust quantitative methods. Both pathogens are members of the order Pleosporales, which also includes pathogens such as *Leptosphaeria* and *Alternaria* spp., and share similar infection strategies ^17^. They each employ necrotrophic effectors that interact with dominant host susceptibility genes to trigger an uncontrolled hypersensitive response, resulting in necrotic lesions that support pathogen proliferation ^18^. Notably, both species share the necrotrophic effector ToxA, with evidence suggesting a horizontal gene transfer event, indicating stable physical proximity between the two species ^19^. Breeding efforts have made significant progress in improving wheat resistance to bothdiseasesthrough the elimination ofdominantsusceptibility genes suchas *TaTsn1*, which interacts with ToxA ^20^, and through the stacking of resistance alleles for both SNB and TS ^21,22^. Despite these advances, both diseases continue to persist in wheat-growing regions worldwide and remain priority targets for resistance breeding ^23^, suggesting that current single-pathogen evaluation approaches may not capture the complexity of field disease dynamics.

Previous efforts to study co-infection dynamics between these pathogens have faced significant methodological constraints. Due to the morphological similarity of lesions produced by both pathogens, and with coalescence obscuring distinct characteristics, visual assessment of disease symptoms requires microscopic analysis even by trained pathologists ^24,25^. Neural-network based disease identification algorithms, which have achieved superior classification accuracy for other wheat diseases, have not yet addressed the challenging differentiation between morphologically similar leaf spot diseases such as SNB and TS ^26^. The development of Real-Time Polymerase Chain Reaction (RT-PCR) assays to quantify both pathogens in host tissue as well as the closely related hemi-biotroph *Zymoseptoria tritici* has provided pioneering insight into the dynamics of foliar wheat pathogens during co infection ^6,27,28^. However, these studies rely on DNAasadirect analogue for pathogen biomasswell as the utilizationof targetswith unstable copy number in the genome including ribosomal and transposon associated sequences. For example, a multicopy target for *P. tritici-repentis* ^29^ which was found to be variable acrossstrainsand regions (Figure S1)^28^. Although this increases its sensitivity, it renders the assay unsuitable for field application where uncharacterised strains may be present. Additionally, RT-PCR suffers from a lack of reproducibility at low pathogens titre, especially in complex environmental samples where PCR inhibitors are known to be present ^30^. Digital PCR (dPCR) addresses limitations of RT-PCRby providing absolute quantification of target nucleic acids without standard curves, offering greater precision and reproducibility while being more tolerant of inhibitors and capable of reliably detecting low abundance targets^31^.

Despite significant advances in breeding for resistance against SNB and TS, both *P. nodorum* and *P. tritici-repentis* continue to persist in wheat-growing regions globally ^6,16,23^. They are most commonly found under co-infection conditions, suggesting that interactions between the two organisms may facilitate their sustained proliferation. Consequently, SNB-TS wheat pathosystem has been a focal point and model of research into polymicrobial infection of plants in recent years, where a range of RT-PCR-based targets have been developed for pathogen monitoring and demonstrated a high field co-infection incidence ^6,16,28^. It is not known how wheatvarieties that differ in resistance to TS and SNBdictate the outcome ofsingle versus SNB-TS co-infection, and so we sought to clarify whether *P. nodorum* and *P. tritici-repentis* form cooperative or competitive associations during co-infection in wheat. We hypothesise that pathogen proliferation will be determined by: (1) genetic resistance present in the host wheat leaves, (2) co-infecting pathogen presence orabsence and (3) pathogenarrivalorder.Here,we develop aduplexdigitalPCR assayalongside automated image analysis enabling simultaneous absolute quantification of both pathogens from infected tissue and apply these tools to investigate polymicrobial disease dynamics in a necrotrophic plant pathosystem.

## Materials and Methods

### Fungal Manipulations

Reference isolates for *P. nodorum* SN15 ^32^ and *P. tritici-repentis* M4 ^33^ were used for all *in vitro* and defined *in planta* inoculation in this study. All isolates were maintained on 90 mm V8 PDA plates ^34^, at ambient temperature under a 12h light/dark photoperiod. To generate mycelial tissue, isolates were grown in Fries 3 broth ^35^beforelyophilisation.To generate inoculum,*P.nodorum*conidiospores were harvested directlyfrom V8PDA platesand resuspended in 0.02% w/v Tween. Inoculum for *P. tritici-repentis* isolates was generated as previously described ^36^.

### Visual Disease Analysis

Harvested leaves were photographed *in situ* using a Google Pixel 6 Pro (USA). Individual leaf images were manually extracted and catalogued using ImageJ ^37^. Differential hue value masks were then generated to distinguish between healthy, chlorotic and necrotic tissue, with cutoffs at 90, 43 and 30 respectively utilizing the opencv library for Python. For full HSV values, refer to Table S1. The Python tools “LeafScan” for image processing and analysis were developed and are available at https://github.com/LeonLenzo/leaf-scan. Normalised pixel values to total leaf areas were used for downstream analyses.

### Nucleic Acid Purification

All fungal and plant tissues were snap frozen in liquid nitrogen and homogenized using a Tissue Lyser II (Retsch, Duesseldorf, Germany), and DNA was purified using the Biosprint DNA Plant Kit in a Biosprint15 (both, Qiagen, Hilden, Germany) DNA concentrations were determined by fluorometric analysis using a Qubit Flex system employing the dsBRDNA Working Solution (Thermofisher, Massachusetts, USA).

### Digital PCR Assay

Primers aT_F and aT_R were designed to simultaneously amplify a 187-bp region in *P. nodorum* and a 185-bp region in *P. tritici-repentis.* These sequences were then used to query 179 *P. nodorum* ^38,39^ and 67 *P. tritici-repentis*^14,40^ genome assemblies toensurewithin-speciesconservationandsingle copy status. Internalfrom the primer-binding-site sequences, species-specific fluorescent probes Pn_FAM and Ptr_HEX were designed to differentially amplify the target organisms (Table 1). Purified gDNA from *P. nodorum* SN15 and *P. tritici-repentis* M4 were used to optimize the assay. dPCR reactions were carried out in 22-µl volumes, where each well contained 11µl ddPCR Supermix for Probes (No dUTP) (Bio-Rad, California, USA), 5μl DNA template, and 0.25µM of primer/probe mix at equimolar concentration. A QX200 Automatic Droplet Generator (Bio-Rad, California, USA) was used to fractionate samples according to manufacturer’s instructions, before undergoing thermal cycling at 95°C for 10 min, then 40 cycles of 94°C for 30 s, 60°C for 60 s, then a final denaturation of 95°C for 10 min. The QX200 Droplet Reader (Bio-Rad, California, USA) was used to quantify positive droplets, with data analysis performed using QuantaSoft Analysis Pro v1.0.596 (Bio-Rad, California) to determine absolute copy numbers following Poisson distribution correction. For downstream analysis, copypermicrolitreoutputs wereonly countedfor values>1, andall input DNA values were normalised to 1ng/μl. In silico off-target analysis was performed by querying all primer and probe sequences against the NCBI RefSeq fungi database using BLAST nucleotide search. BLAST results were filtered using minimum sequence identity thresholds of 60% for primersand 85% forprobes. Cross-reactive organisms were identified as those showing significant hits to both primers(aT_Fand aT_R) and at least one species-specific probe (Pn_FAM or Ptr_HEX), representing potential targets for non-specific dPCR amplification (Figure 1D). Primers and probe specificity were tested *in vitro* using the ddPCR assay (Figure 2C) against the purified gDNA of closely related fungal pathogens and wheat (Table S2).

**Figure 1:**
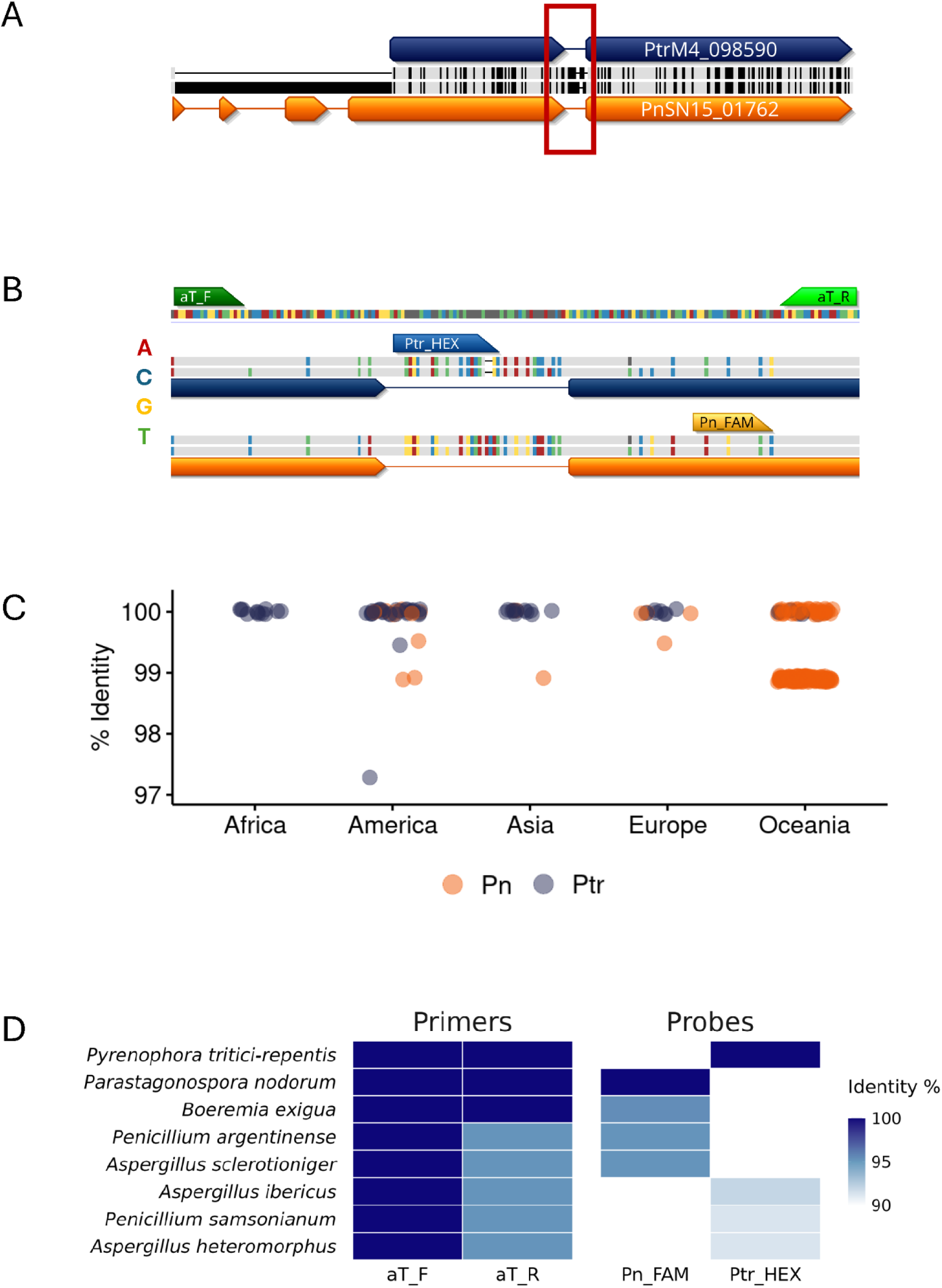
dPCRDesign **(A)** Pairwise alignment of PnSN15_01762 (Orange) and PtrM4_098590 (Blue), the orthologues for alpha tubulin. Locus aT is highlighted in red. **(B)** Primer and probe design showing aT_F and aT_R amplification primers (green arrows) with species-specific probes Ptr_HEX (blue) and Pn_FAM (orange). Coloured bars indicate sequence polymorphisms. **(C)** Percentage Identity of the aT region across pan-genome assemblies. **(D)** Heatmap showing sequence identity between primer/probe sequences and organisms in the Refseq fungi database.

**Figure 2:**
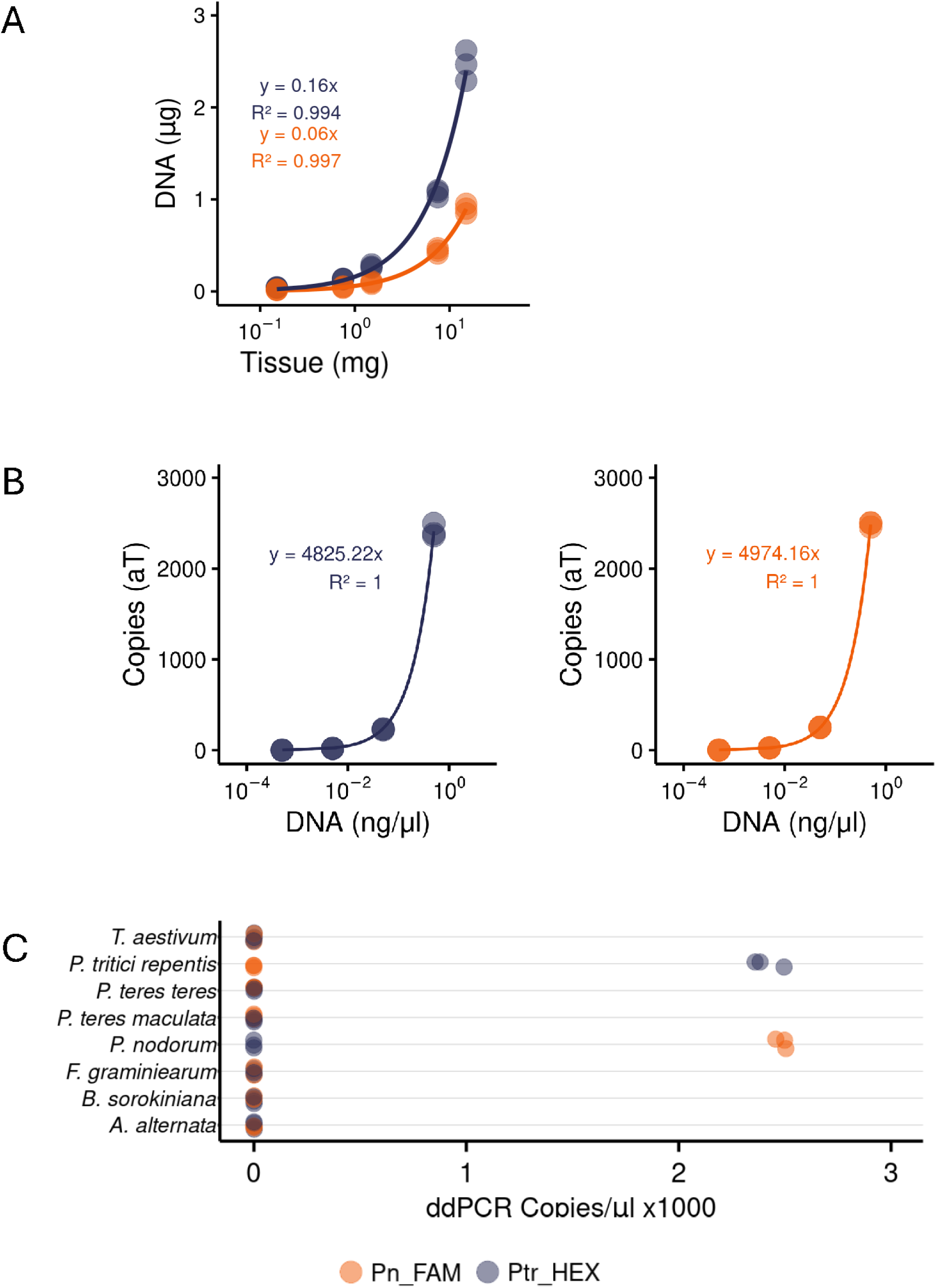
dPCRValidation **(A)**Standard curves showing linear relationship (β) between DNA yield for *P. tritici-repentis* (blue) and *P. nodorum* (orange). **(B)** Standard curves for Ptr_HEX (blue) and Pn_FAM (orange) probes showing linear relationship (α) between DNA concentration and copy number. **(C)** Specificity testing of the dPCR assay against gDNA from wheat host and related fungal species, showing probe-specific amplification patterns.

**Table 1.**
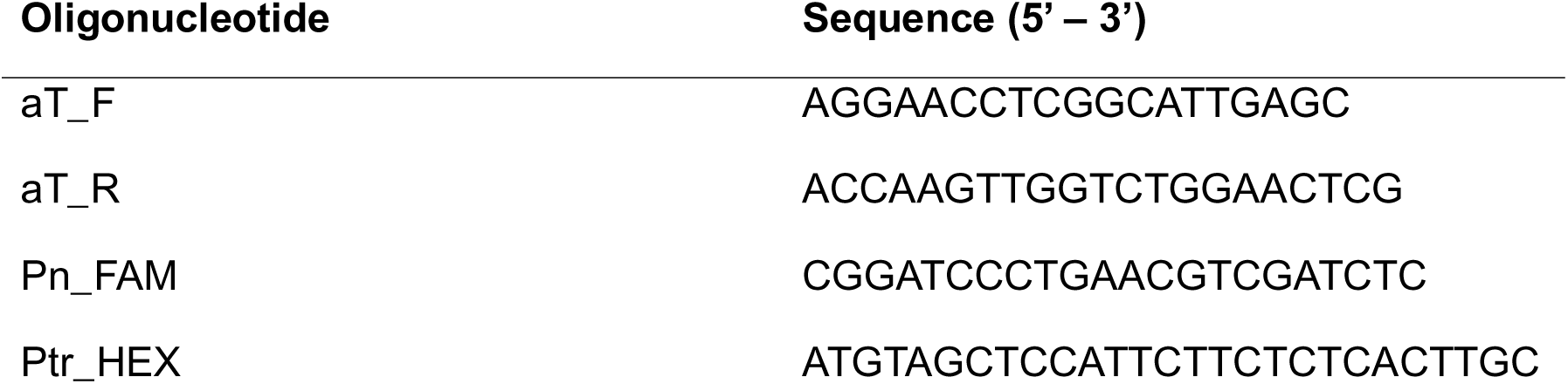
Oligonucleotides used in this study.

### Biomass Modelling

To approximate a relationship between dPCR copy number and fungal biomass, we developed a two-step calibration approach. First, wequantified the DNAyieldper unit fungal tissue byextracting DNA from known quantities of freeze-dried mycelium. Three biological replicates of *P. nodorum* SN15 and *P. tritici-repentis* M4 mycelia were weighed out in technical triplicates between 0.15 and 15 mg and DNA was extracted as described above. Total DNA content was plotted as a function of tissue mass and analysed using linear regression to derive the species-specific DNA-to-tissue ratio (β) for each pathogen. Second, we established the relationship between DNA input and dPCR target copy number by analysing serial dilutions of gDNA from both species. Triplicate samples at concentrations ranging from 0.05 to 500 pg/µl were analysed using the dPCR assay described above. The integrated biomass model combines these relationships through the formula below where C represents dPCR-detected alpha-tubulin copies/μl, α is the target copies per nanogram of DNA, and β is the micrograms of DNA per milligram of tissue. After substituting the experimentally derived values, we obtained simplified predictive equations for each species that enable direct conversion of dPCR-measured target copies into estimated fungal biomass.

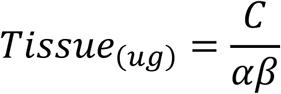

### Field Sites and Experimental Design

Symptomatic leaf tissue wascollected between May and Octoberof 2022 from field sites sown in early April in Western Australia. The first, a disease nursery unattenuated by fungicide for more than ten years (Northam, 31°32’23.4“S 116°42’17.3”E). The second plot (South Perth,-31.991046, 115.888172), contained three adjacent plots, established by incorporating infected wheat stubble containing single or mixed pathogen species into the soil prior to sowing, with cross-contamination between adjacent plots expected over the growing season to test competitive dynamics under varying establishment priorities (Figure S2). Cultivars were selected for resistance through Department of Primary Industries and Regional Development of Western Australia wheat variety disease ratings to select lines with differential susceptibility to SNB and TS (Table 2). SVS (susceptible-very susceptible) indicates wheat cultivars with minimal disease resistance, while MRMS (Moderately Resistant-Moderately Susceptible) denotes cultivars with intermediate resistance levels. Samples were photographed on site, trimmed with a scalpel to 50 mm in length, then stored on ice for ≤ 24h before snap freezing and storage at-80°C until processing.

**Table 2.**
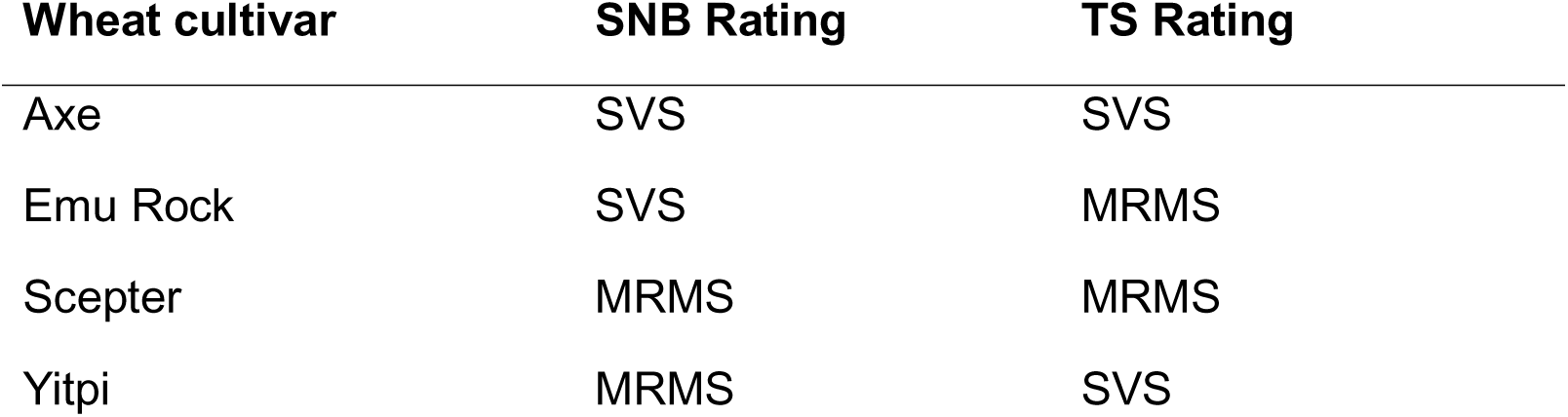
Australian wheat cultivars used in this study and their disease resistance ratings ^64,65^. SVS, susceptible-very susceptible; MRMS, moderately resistant-moderately susceptible.

### Attached Leaf Assay

Wheat seedlings were grown under white light on a 12hr photoperiod until 12 days old. Leaves were mountedadaxialside uponto 45mm XPSfoamboardsusing micropore tapeinside aseedling tray. Attached leaveswere inoculated with 15µlofeither 1×10^7^spores.ml^-1^ of *P. nodorum*SN15 spores, 1×10^5^spores.ml^-1^ of *P. tritici-repentis* M4 spores suspended in 0.02% Tween 20 or a mixture of the two maintaining the original concentration. Following inoculation spore suspension was spread with a paintbrush (WestArt, 4mm Synthetic), leavesallowed todry, then sealed inside humidity chambers and maintained at 22C± 1C, ≥ 95% humidity under white light on a 12 hr photoperiod. Staggered infections were carried out 72 hrs post-initial inoculationfollowing the sameprocedure. Leaveswerephotographedsevendaysafter secondary infection, before excising a 50mm section with a sterile scalpel.

## Statistical analysis

Statistical analyses wereperformed using Rversion 4.5.0 ^41^. Linear regression analysiswasconducted using the lm() function to fit a linear model. Pathogen biomass data were log10-transformed to improve normality and stabilize variance. Normality was assessed using Shapiro-Wilk tests, which confirmed that log10 transformation substantially improved the distribution of biomass data. Student’s t-tests were used to compare pathogen biomass between co-infection and single infection treatments within each site and wheat line. For sequential inoculation data, one-way ANOVA was conducted separately for each wheat line with treatment as the factor. Post-hoc comparisons used Tukey’s HSD test when significant effects were detected (p < 0.05). ANOVA assumptions were verified through residual analysis and Shapiro-Wilk tests for normality.

## Results

### Biomassmodelling and absolute pathogen DNAquantification using dPCR

To identify a suitable locus for marker development, gene annotations from SN15 and M4 were compared using reciprocal BLASTanalysistoidentify highly conserved orthologues. Candidates were filtered manually to identify regions ≤200 bp with conserved 5’ and 3’ termini for primer binding sites adjacent to a divergent central region for a species-specific probe, before selecting a region spanning the final intron of the alpha-tubulin ortholog in both species (Pn_SN15_01762, Ptr_M4_98590). Suitable loci within the orthologous alpha-tubulin coding sequences were identified so that a single primer set could target binding sites conserved across both species, while species-specific internal probes (Pn_FAM and Ptr_HEXrespectively) permitted species discrimination (Figure 1A & B). Single copyprimer-probe sites were conserved at a single copy across pan-genomes of the respective species, with in silico cross-reactivity analysis confirming minimal risk of in species off target effects(Figure 1C). No cross reactivity to known wheat associated fungi was observed when tested in silico, with only seven primer +probe matches≤ 98% found excluding target organisms (Figure 1D). Species-specific DNA yield analysis revealed *P. tritici-repentis* produces 0.160 μg DNA per mg dry tissue compared to *P. nodorum* at 0.059 μg DNA per mg dry tissue (Figure 2A). Calibration curves for both probes demonstrated near-perfect linearity (R² > 0.99) across a broad range of fungal DNA (0.5 pg.μl^-1^ to 0.5 ng.μl^-1^) for both pathogens (Figure 2B). When tested against gDNA of the wheat host or related fungal species and alternative wheat pathogens, the assay showed no off-target effects (Figure 2C). The integrated biomass model yielded conversion factors of 1.3 μg biomass per target copy for *P. tritici-repentis*and 3.4 μg per target copy for *P. nodorum*, providing thefoundation for allsubsequent comparative analyses (Figure S3).

### Incidence ofhost tissuenecrosis(but not chlorosis) correlates with fungal biomass accumulation

Total fungal biomass measured through dPCR correlated with necrosis measured with automatic pixel thresholding (Figure 3A). Controlled infection of wheat cvs Axe and Scepter with single or mixed *P. nodorum*/*P. tritici-repentis* inoculum were monitored through photography and tissue sampling over a 10-day post-infection period. Linear regression models were constructed for each of three distinct disease metrics: chlorosis, necrosis and total disease (necrosis plus chlorosis) (Figure 3B). Of the three metrics evaluated, necrosis demonstrated the strongest positive correlation (R^2^= 0.705). Total disease coverage (chlorosis plus necrosis) exhibited a moderate positive correlation with pathogen biomass (R^2^= 0.445). Interestingly, chlorosis showed a weak negative correlation with pathogen biomass (R^2^= 0.214), suggesting that chlorotic symptoms are not indicative of high pathogen loads. We then extended this analysis to naturally infected wheat plants sampled weekly over a 16-week period beginning at tillering stage at a field site. Field data revealed substantial temporal variation in disease-biomass correlations (Figure 3C). Strong correlations similar to controlled conditions were observed in week 1, but collapsed and recovered across the season, likely reflecting temporal confounding as infections at varying stages are sampled simultaneously. This temporal pattern highlighted the importance of assessment timing for the correlation between visual symptom assessment and fungalbiomass.

**Figure 3:**
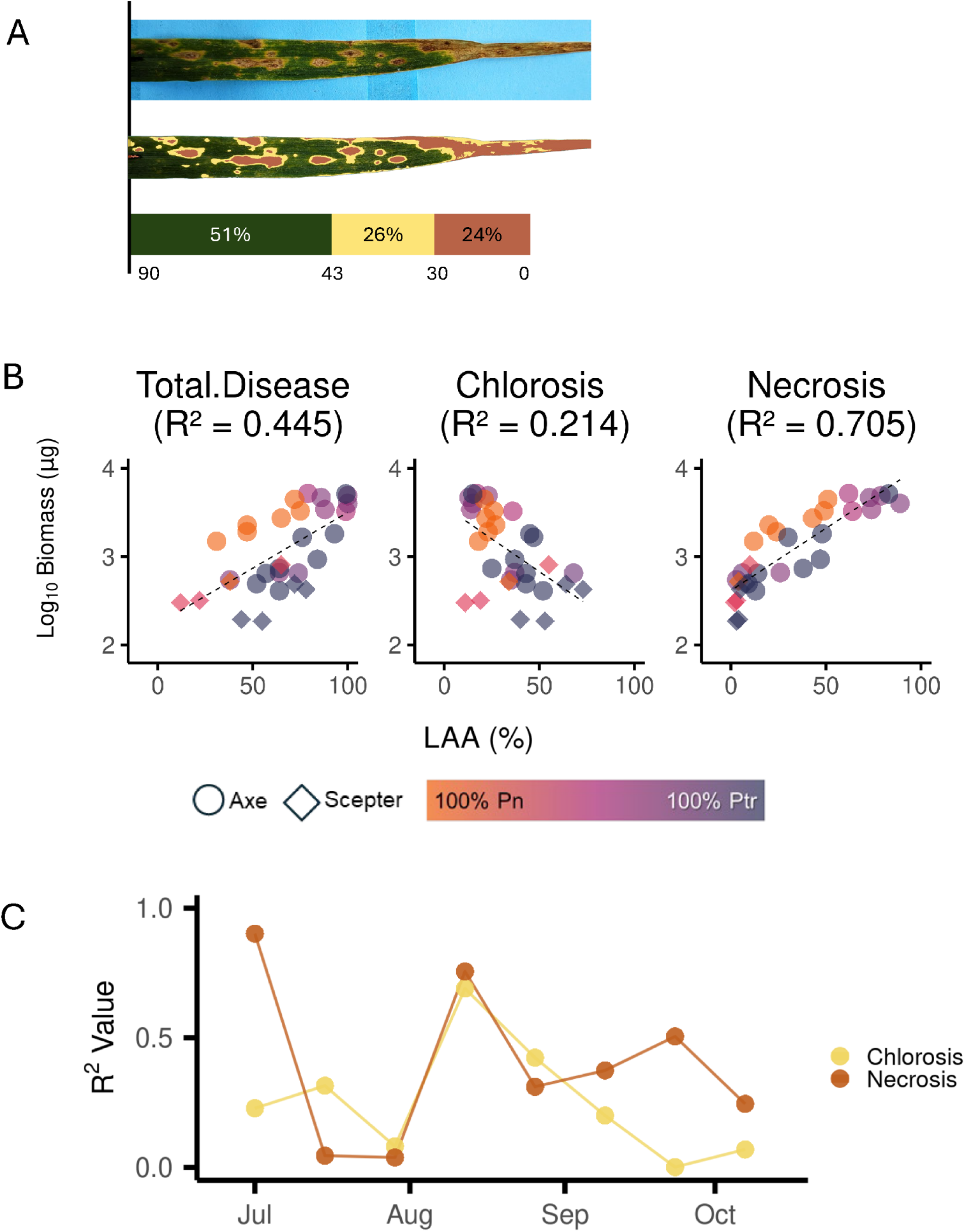
Diseasebiomasscorrelation **(A)** Automated leaf segmentation showing original image, processed output with disease classification (green = healthy, yellow = chlorotic, brown = necrotic), and HSV hue thresholds. **(B)** Correlations between pathogen biomass and leaf area affected (LAA) by different visual disease metrics. Points represent Axe (susceptible, circles) and Scepter (resistant, diamonds) varieties with single or mixed infections. Colour gradient indicates *P. nodorum* (orange) to *P. tritici-repentis* (blue) proportion. R² shows correlation strength for total disease, r shows correlation co-efficient. **(C)** Temporal variation in visual disease-biomass correlations under field conditions over a 16-week sampling period. Lines show R² values for chlorosis (yellow), necrosis (brown) metrics.

### Fieldco-infection facilitates biomass accumulation and the breakdown of host resistance

Pathogenbiomassresponsesto geneticresistance in host plants were examinedinfour wheat cultivarswith combinatorial differences in TS and SNB disease resistance ratings (Table 1), sampled in a naturally wheat-disease nursery endemic for both *P. nodorum* and *P. tritici-repentis*. Co-infected samples were the most common type observed (63%) validating previous observations ^6,16^, with single infections representing 21% and 16%of samplesfor *P.tritici-repentis* and *P.nodorum,*respectively(Figure 4). For Axe (SVS to both TS and SNB), no significant differences in necrosis severity wereobserved between co-infected and single-infected samples, with significantly higher *P. tritici-repentis* but not *P. nodorum* biomass observed under co-infection. With Emu Rock and Yitpi, the only significant difference observed was a higher proportion of *P. nodorum* biomass in Emu Rock, with no single infections of a given pathogen observed in hosts with a resistance to said pathogen.

**Figure 4:**
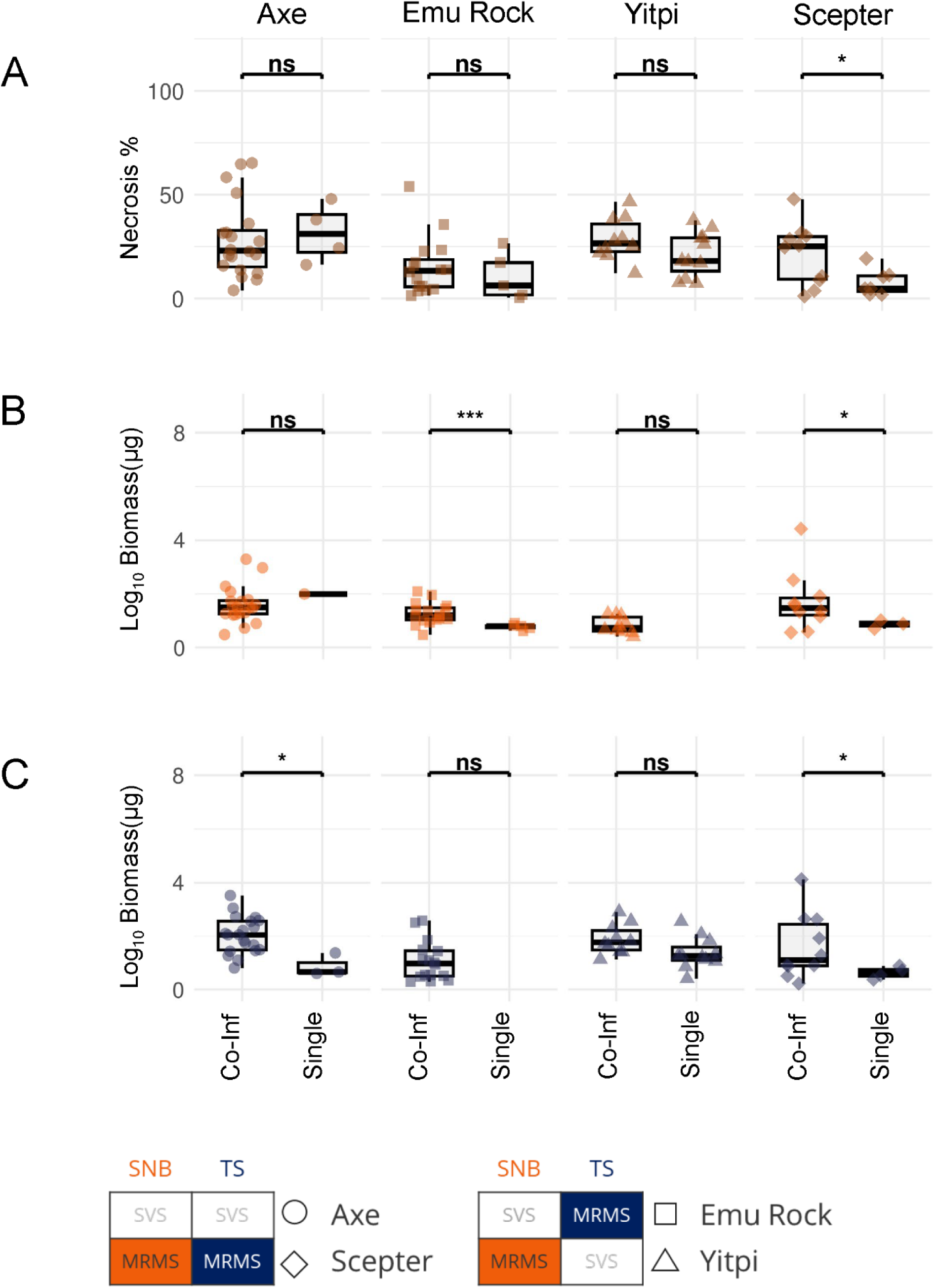
Host genetics innatural field disease **(A)** disease severity (necrosis %, brown), **(B)** *P. nodorum* biomass (orange), and **(C)** *P. tritici-repentis* (blue) biomass across four wheat lines: Axe (SVS to SNB and TS), Emu Rock (SVS to SNB, MRMS to TS), Scepter (MRMS to SNB and TS), and Yitpi (MRMS to SNB SVS to TS). Points represent individual wheat leaves with shapes indicating cultivar (circles = Axe, squares = Emu, diamonds = Scepter, triangles = Yitpi). Statistical significance of differences between co-infection and single infection within each wheat line indicated above brackets (ns = not significant, * p < 0.05, ** p < 0.01, *** p < 0.001).

Wheat cv. Scepter, which is rated MRMS to both TS and SNB, demonstrated greater necrosis when both pathogens were present (Figure 4A). Both *P. tritici-repentis*and *P. nodorum*were detected atvery lowsingle-infection frequencies and significantly less biomass than co-infected samples. In general, both pathogens exhibited significantly higher biomass accumulation under co-infection conditions compared to single-pathogen infections (Figures4B & C). Notably, the only wheatcultivar toexhibit higher necrosis and biomass during co-infection was Scepter, the host with the highest resistance toboth pathogens.

### Fair weather friends: P. tritici-repentisfacilitates co-infection while P. nodorumsuppresses

We sought to isolate priority effects in the field by using a defined primary inoculum in a field trial involving three adjacent plots, capable of spreading airborne inoculum for secondary infections. Exterior plots containedeitherstubble infected with*P.tritici-repentis*or *P.nodorum*and the centralplot containedstubble with an equal mixture of both (Figure S2). For the *P. tritici-repentis* and mixed stubble site, co-infection was again ubiquitous throughout the growing season, suggesting the presence of secondary airborne inoculum^7^. Disease incidences measured by necrosis and biomass for both species were significantly higher under co-infection than for single infection, with the exception of *P. nodorum* at the mixed site, broadly reflecting the patterns seen under endemic conditions (Figures 5, S4 and S5).

**Figure 5:**
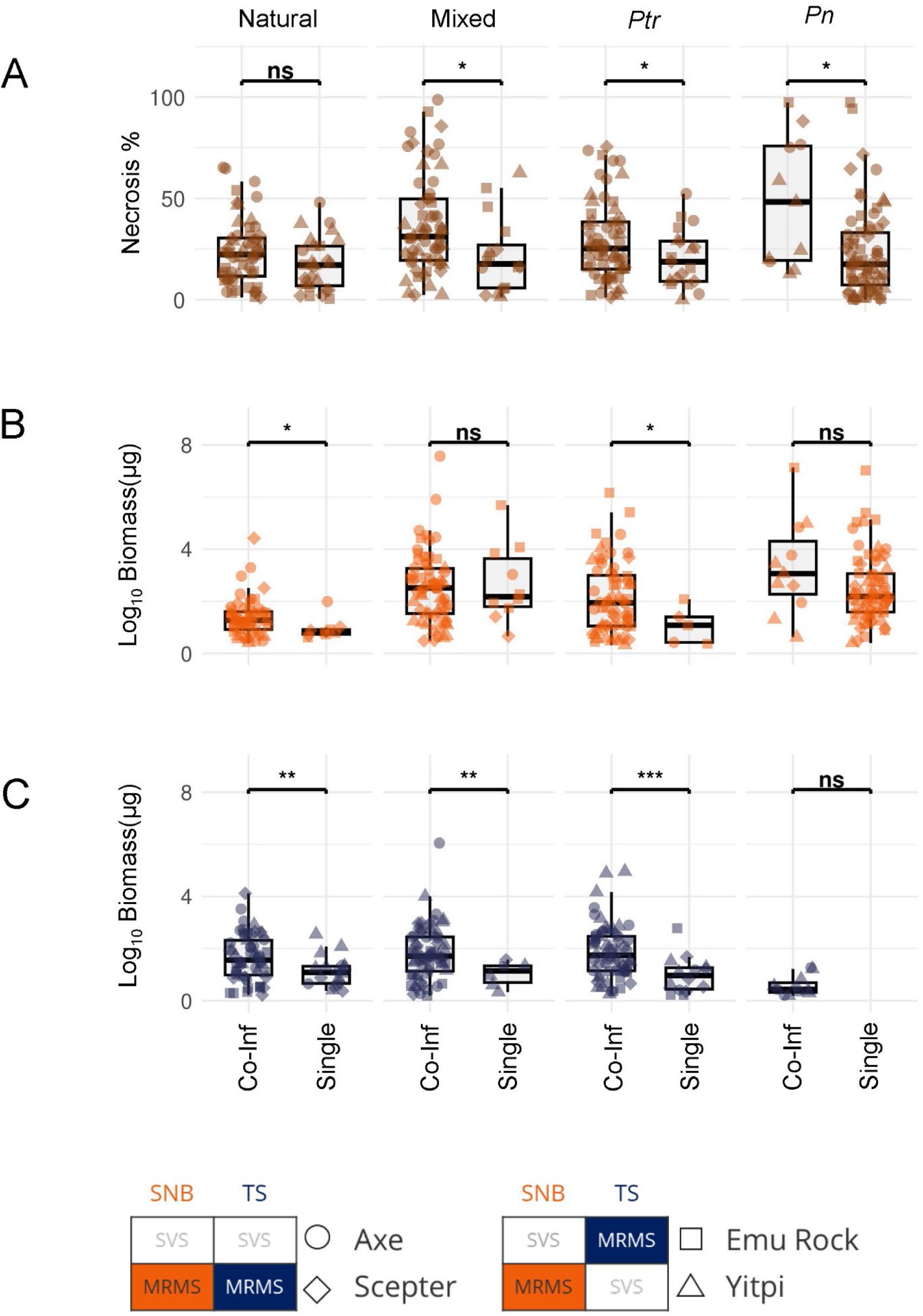
Natural vs defined field inoculum **(A)** necrosis severity (necrosis %, brown), **(B)** *P. nodorum* biomass (orange), and **(C)** *P. tritici-repentis* biomass (blue) across four field sites. Natural (naturally occurring infection in an endemic field Figure 4), Mixed (equal mix of *P. nodorum* and *P. tritici-repentis*infected stubble), *Pn (P. nodorum* infected stubble) and *Ptr* (*P. tritici-repentis* infected stubble). Points represent wheat leaf samples with shapes indicating wheat lines (circles = Axe, squares = Emu, diamonds = Scepter, triangles = Yitpi). Statistical significance of differences between co-infection and single infection within each site indicated above brackets (ns = not significant, * p < 0.05, ** p < 0.01, *** p < 0.001).

The *P. nodorum* inoculum site revealed dramatically different interaction dynamics. *P. tritici-repentis*failed to establish single-pathogen infection and was only observed at a low frequency in co-infected samples (Figures5 and S6). Thissuggests that *P.nodorum*may antagonise secondary *P.tritici-repentis* host-infection indicating priority effects in the tripartite *P. nodorum*-*P. tritici-repentis*-wheat.

### Controlled sequential infection reveals an asymmetric partnership

Toconfirmthisantagonism,we conducted sequentialco-infection assaysto investigate the roleoftemporal dynamics. *P. nodorum* and *P. tritici-repentis*were inoculated onto wheat cvs. Axe and Scepter seedlings in acontrolled environmental growth chamber, wheresecondary infection wascarried out 72hrsafter primary infection using defined pathogen isolates. Analysis of wheat cv. Axe revealed consistent biomass accumulation and necrosis development across treatments as well as secondary-infection timings (Figure 6A). Simultaneous inoculation resulted in *P. tritici-repentis* dominance, while *P. tritici-repentis* priority led to balancedpathogen loadswith nosignificant competitive advantage for either species pertaining to biomass accumulation. When *P. nodorum* was applied as a primary inoculum ahead of *P. tritici-repentis*, it again inhibited subsequent *P. tritici-repentis* biomass accumulation. This was the strongest competitive asymmetry observed across all treatments on wheat cv. Axe and provides additional evidence of the inhibitory dynamics observed under field conditions.

**Figure 6:**
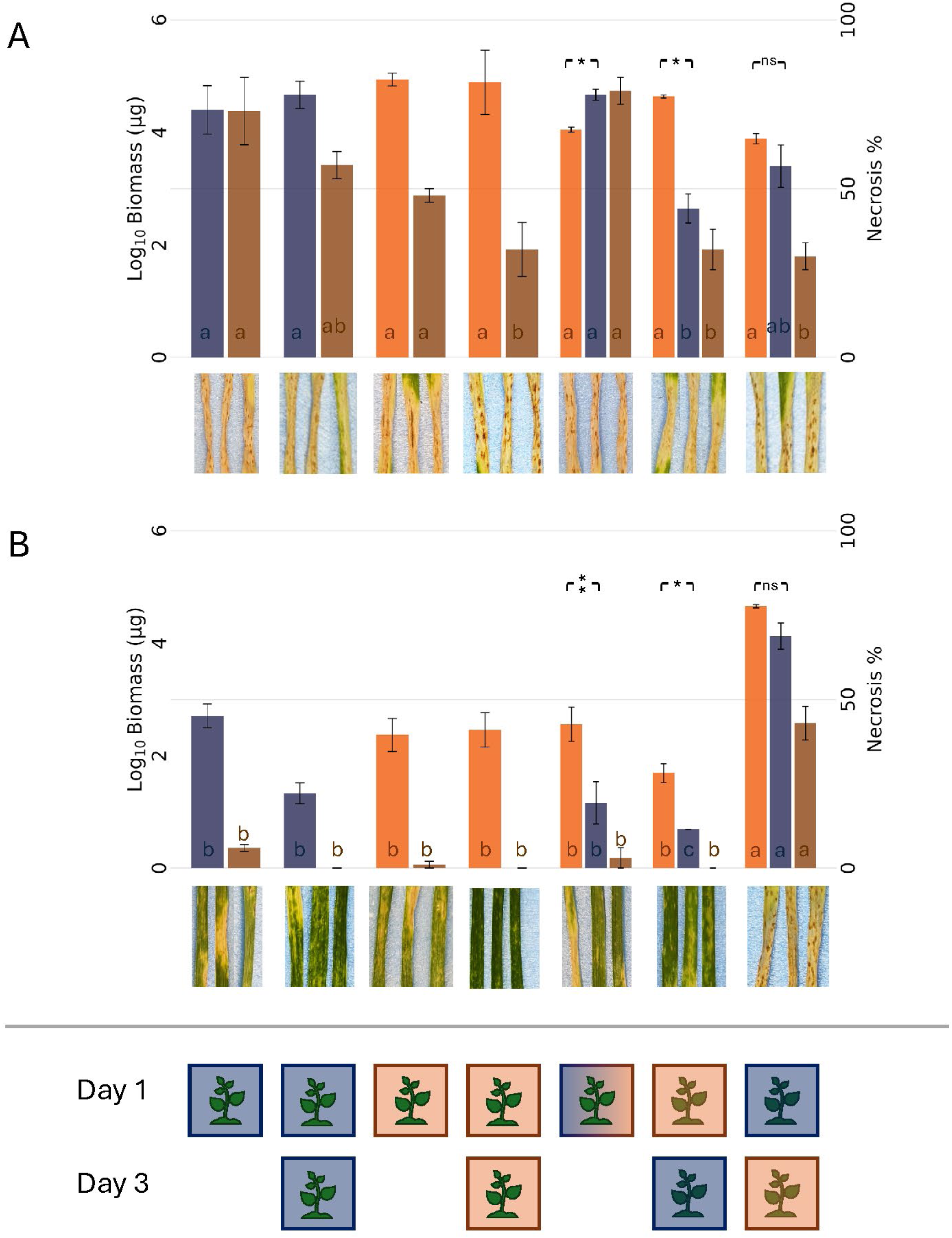
Sequential Seedling Infection **(A)** Axe and **(B)** Scepter. Bars represent mean (±SE) log₁₀ biomass (μg) for *Pn* (orange) and *Ptr* (blue), with Necrosis % (brown) plotted on the secondary y-axis. From left to right, Single pathogen treatments: Ptr → Ø Ptr → Ptr, Pn → Ø, Pn → Pn, Co-infection treatments: Pn + Ptr, Pn → Ptr, Ptr → Pn. Letters inside bars indicate statistical groupings within each series (Tukey HSD, α = 0.05). Asterisks above bars indicate significant differences between *Pn* and *Ptr* biomass within co-infection treatments (paired t-test: ns = p ≥ 0.05, * = p < 0.05, ** =p < 0.01,). n = 3 per treatment.

Sequential co-infection on wheat cv. Scepter displayed markedly different infection dynamics. Most treatments resulted in substantially reduced pathogen biomass and necrosis compared to wheat cv Axe, likely due to background resistance (Figure 6B). *P. tritici-repentis* followed by *P. nodorum* inoculation was the sole exception, representing a breakdown of seedling resistance where both pathogens achieved biomass levels comparable to those in susceptible plants. This treatment was also the only one to induce significant necrosis in resistant plants, suggesting that *P. tritici-repentis* establishment compromises subsequent resistance responses to *P. nodorum,* allow both pathogens to proliferate.

Collectively, this work confirms priority effects influence pathogen proliferation in the SNB-TS/wheat pathosystem. Under natural field conditions, co-infection is ubiquitous and facilitates biomass accumulation for both pathogens, synergistically break down host resistance. When the inoculum source is controlled, *P. nodorum* suppresses *P. tritici-repentis* establishment, while *P. tritici-repentis* facilitates balanced co-infection. In controlled sequential infections, *P. tritici-repentis* primacy enables both pathogens to overcome genetic resistance, while *P. nodorum* primary infection consistently suppresses *P. tritici-repentis* infection regardless of host background.

## Discussion

Co-infection dynamics play crucial roles in wheat pathosystems, affecting pathogen virulence and epidemiology. Priority effects have been shown to impact disease development and competitive advantage ^43^, immune suppression enabling persistence ofnormallyavirulent strains ^44^, andeffector complementation betweenstrains enhancing disease severity ^45^. The challenge ofaccurately measuring pathogenabundance in polymicrobial leaf spot diseases has been well established prior to this study ^24,25^. Previous studies using RT-PCR were able to highlight the high frequency of wheat co-infection by *P. nodorum* and *P. tritici-repentis*, however the work was limited by available technology to measuring relative pathogen abundance ^6,27,28^.

Our methods represent a significant advancement in the study of polymicrobial plant diseases. Through species-specific DNA yield normalization, we showed *P. tritici-repentis*yielded almost three times the DNA per milligram of tissue compared to *P. nodorum*. Using a conserved single copy gene eliminates variability between strains while the utilization of dPCR retainssufficient sensitivity for field applications. This provided a strong foundation to investigate the relationship pathogen abundance and visual symptoms, wheat cultivar disease resistance ratings, and the role of priority effects in the outcome of disease. Our findings reveal two interactions. The first is co-operative, where *P. tritici-repentis* priority establishment facilitates subsequent *P. nodorum* colonisation and enhances disease severity and biomass for both pathogens in a resistant host. The second is competitive, where *P. nodorum* priority establishment suppresses *P. tritici-repentis* colonisation regardless of host resistance background. To our knowledge, this is the first study to observe asymmetricfacilitation overcoming host resistance in a fungal plant pathosystem.

Fieldtrials where *P. tritici-repentis*wasintroduced first reproduced facilitation patternsobserved innaturally infected fields. Priority establishment by *P. tritici-repentis* may drive the co-infection observed in wheat production systems, where *P. tritici-repentis* ascospores have been shown to mature up to three months before those of *P. nodorum* in south-west Western Australia ^46,47^. In addition, *P. tritici-repentis* may benefit from superior establishment characteristics, including larger resource-rich and temperature-resistant conidia ^48,49^.When *P.tritici-repentis*establisheson resistant host leavesit facilitates *P.nodorum*colonisation leading togreater diseasethrough anunknown mechanism thatleadspositive interferencewhere temporal sequence determines resistance breakdown. *Zymoseptoria tritici* followed by *Pseudomonas syringae* infection leads to systemically induced bacterial susceptibility in wheat, where fungal infection facilitates colonization through suppression of immune-related metabolites ^50^. Priority effects may exploit depleted immune responses, where initial infection activates energetically costly defence signalling pathways creating windows of vulnerability for secondary infections ^51^. Necrotrophic pathogens can subvert plant defence responsesthat evolvedprimarily against biotrophic pathogensincluding thesalicylicacid pathway, which while providing resistance against biotrophs, can enhance susceptibility to necrotrophs ^52^.

Necrotrophic pathogens create favourable conditions for subsequent colonisation by generating necrotic lesions with effectors that cause necrosis and provide nutrient-rich substrates from lysed plant cells, support necrotrophic growth ^53^. Our observations are distinguished by the asymmetric nature of this facilitation despite both pathogens producing necrotrophic effectors infection on wheat ^14,39^. This suggests that mechanistic differences beyond necrotrophic virulence determine the facilitative outcomes we observed. Previously, *P. nodorum* strains producing high levels of necrotrophiceffectors rescued reduced virulence strainson susceptible wheat,maintaining overall virulence of the pathogen population ^45^. Similarly the introduction reduced virulence mutants of *Magnaporthe oryzae* into populations of wild-type strains resulted in enhanced overall population fitness and increased damage to rice hosts ^11^. These examples illustrate that cooperation between pathogen strains with different virulence capabilities can lead to enhanced disease outcomes, representing a distinct mechanism from host immune suppression-mediated facilitation. The mechanism underlying the facilitation observed in our data remains unclear, but could involve direct pathogen-pathogen interactions,host immunesuppression,or bothprocesses working in combination ^9^.

*P. nodorum* priority establishment antagonises *P. tritici-repentis* colonisation across all host genotypes observed in the field and laboratory, suggesting a physiological mechanism possessed by *P. nodorum* to suppress the growth of competitors. While *P. nodorum* and *P. tritici-repentis* share a near-identical ToxA ^19^, *P. nodorum* possesses a diverse collection of characterised necrotrophic effectors including SnTox1, SnTox3, SnTox5 ^54^and SnTox267 ^55^. *Verticillium dahliae*has co-opted antimicrobial proteins specifically for manipulating fungal competitors during host colonization ^56^, suggesting exclusion through biochemical interference may be an evolved strategy rather than an incidental consequence of infection. It is not known if characterised *P. nodorum*NEs possess antimicrobial functionshowever, SnTox1 possessdual functions; facilitatingnecrosisin wheat and bindingchitin,possibly to evade host PAMP-triggeredrecognition ^57^.Recent pan-genomic analysis of *P. nodorum* identified an abundance of candidate secreted effector proteins, including numerous homologs to known antimicrobial effectors from other fungal species ^38^. In addition, *P. nodorum* possesses an extensive arsenal of secondary metabolite biosynthesis genes ^58^. Previous studies revealed that *P. nodorum*was capable of producing volatile organic compounds that exhibit phytotoxic and antimicrobial activities ^49,59,60^. It remains to be seen if these secondary metabolites possess bioactivity that antagonises *P. tritici-repentis* ^61^.

Molecular quantification revealed competitive exclusion patterns masked by visual assessment. While necrosis provides a reliable estimate of total pathogen biomass under controlled conditions, visual symptoms cannot reliably distinguish between single and co-infection states.Additionally, quantification of disease can be hindered by simultaneous biotic and abiotic stresses, emphasising the need for molecular approaches that quantify pathogens biomass ^62^. Even with automated image analysis tools like LeafScan, which can accurately quantify necrotic tissue area, morphologically indistinguishable lesions produced by *P. nodorum*and *P. tritici-repentis* provide novisual indication of their microbial composition. The correlation observed under controlled infection suggests that both pathogens primarily colonise necrotic tissue regions, likely secreting necrotrophic effectors into surrounding tissue to induce chlorosis ahead of colonisation, supporting previous mechanistic studies ^63^.

These sequence-dependent interactions represent a form of historical contingency where early colonisers fundamentally alter the trajectory of microbial community development within host tissues ^1^. Thefacilitative and competitive mechanisms we observed demonstrate that pathogen communities are not simply assembled through independent colonisation events, but rather through complex ecological networks where arrival order determines interaction outcomes ^5^. This challenges traditional views of agricultural pathosystems as simplified host-pathogen dyads and instead positions them as dynamic microbial ecosystemsgoverned bycommunity assembly rules similar to those operating innatural systems^2^, butwith accelerated temporal dynamics due to the intensive selection pressures of agricultural environments ^3^. Agricultural monocultures may thus serve as model systems for understanding rapid microbialcommunity assembly processes under simplified environmental conditions, offering insights into fundamental ecological principles that operate across diverse ecosystems while providing practical applications for sustainable disease management strategies.

Severalprioritiesemerge from thiswork. Werecommend thatfuturedisease resistance trialsof commercial wheat cultivars consider rating for the SNB-TS complex over the current single disease resistance rating system in wheat-growing regions where *P. nodorum* and *P. tritici-repentis* are endemic. Furthermore, additionalanalysis of the effect that co-infection mayhaveongrain yields is needed. Field validation studies across diverse environments and wheat genetic backgrounds are needed to determine the generalisability of our sequence-dependent findings, particularly investigating whether similar priority effects occur in cultivars with different resistance backgrounds.

## Acknowledgements

The authors would like to thank; Noel Knight (Curtin University) and Lincoln Harper (Curtin University) for assistance with dPCR assay design; Julie Lawrence (Curtin University) and Manisha Shankar (DPIRD) for providing off-target isolatesfor dPCRvalidation.We extend our thanks to Dr Araz Abdullah(Curtin University) for insightful discussions and for establishing the precursor groundwork for this study. This study was conducted by the Centre for Crop and Disease Management, a co-investment between the Grains Research and Development Corporation (GRDC) and Curtin University - grant CUR1403-002BLX.

## Data Accessibility Statement

Alldata supporting the results ofthisstudy are availableasfollows: RawdPCRdata, imageanalysisoutputs, and statistical analysis code are archived in the Dryad Digital Repository at [https://doi.org/10.5061/dryad.mcvdnckch]. The LeafScan image processing and analysis tools are freely available at https://github.com/LeonLenzo/leaf-scan. Reference genome assemblies for *Parastagonospora nodorum* SN15 and *Pyrenophora tritici-repentis* M4 are publicly available through NCBI GenBank. Primer and probe sequences for the duplex dPCR assay are provided in the Materials and Methods section. Field sampling coordinates and experimental metadata are included in the supplementary materials. All materials and protocols are available upon reasonable request to the corresponding author.

## References

1. Fukami T. Historical Contingency in Community Assembly: Integrating Niches, Species Pools, and Priority Effects. Annu. Rev. Ecol. Evol. Syst. 2015 46:1–23. 10.1146/annurev-ecolsys-110411-160340.

2. Vacher C, A Hampe, AJ Porté, et al. The Phyllosphere: Microbial Jungle at the Plant-Climate Interface. Annu. Rev. Ecol. Evol. Syst. 2016 47:1–24. 10.1146/annurev-ecolsys-121415-032238.

3. McDonald BA, EH Stukenbrock. Rapid emergence of pathogens in agro-ecosystems: global threats to agricultural sustainability and food security. Philos. Trans. R. Soc. B Biol. Sci. 2016 371. 10.1098/rstb.2016.0026.

4. Halliday FW, RM Penczykowski, B Barrès, et al. Facilitative priority effects drive parasite assembly under coinfection. *Nat*. Ecol. Evol. 2020 4:1510–21. 10.1038/s41559-020-01289-9.

5. Debray R, RA Herbert, AL Jaffe, et al. Priority effects in microbiome assembly. Nat. Rev. Microbiol. 2022 20:109–21. 10.1038/s41579-021-00604-w.

6. Fejer Justesen A, B Corsi, A Ficke, et al. Hidden in plain sight: a molecular field survey of three wheat leaf blotchfungaldiseasesin North-Western Europe showsco-infection iswidespread. Eur.J. Plant Pathol. 2021 160:949–62. 10.1007/S10658-021-02298-5.

7. Vander Heyden H, P Dutilleul, J-B Charron, et al. Monitoring airborneinoculumforimproved plantdisease management. A review. Agron. Sustain. Dev. 2021 41:40–40. 10.1007/s13593-021-00694-z.

8. Ferreira RBR, LCM Antunes, N Sal-Man. Pathogen-pathogen interactions during co-infections. ISME J. 2025 19:wraf104. 10.1093/ismejo/wraf104.

9. Dutt A, D Andrivon, CLe May. Multi-infections, competitive interactions, and pathogen coexistence. Plant Pathol. 2022 71:5–22. 10.1111/ppa.13469.

10. Harrison F, LE Browning, M Vos, et al. Cooperation and virulence in acute Pseudomonas aeruginosa infections. BMC Biol. 2006 4:21. 10.1186/1741-7007-4-21.

11. Lindsay RJ, MJ Kershaw, BJ Pawlowska, et al. Harbouring public good mutants within a pathogen population can increase both fitness and virulence. eLife 2016 5:e18678. 10.7554/eLife.18678.

12. Mousa WK, A Schwan, J Davidson, et al. An endophytic fungus isolated from finger millet (Eleusine coracana) produces anti-fungal natural products. Front. Microbiol. 2015 6:1157. 10.3389/fmicb.2015.01157.

13. Orton ES, JKM Brown. Reduction of growth and reproduction of the biotrophic fungus Blumeria graminis in the presence of a necrotrophic pathogen. Front. Plant Sci. 2016 7:742–742. 10.3389/FPLS.2016.00742/BIBTEX.

14. Gourlie R, M McDonald, M Hafez, et al. The pangenome of the wheat pathogen Pyrenophora tritici-repentis reveals novel transposons associated with necrotrophic effectors ToxA and ToxB. BMC Biol. 2022 20. 10.1186/s12915-022-01433-w.

15. McDonald BA. How knowledge of pathogen population biology informs management of Septoria nodorum blotch on wheat. Eur. J. Plant Pathol. 2025 171:531–45. 10.1007/s10658-024-02996-w.

16. Abdullah AS, MR Gibberd, J Hamblin. Co-infection of wheat by Pyrenophora tritici-repentis and Parastagonospora nodorum in the wheatbelt of Western Australia. Crop Pasture Sci. 2020 71:119–27. 10.1071/CP19412.

17. Möller M, EH Stukenbrock. Evolution and genome architecture in fungal plant pathogens. Nat. Rev. Microbiol. 2017 15:756–71. 10.1038/nrmicro.2017.76.

18. McDonald MC, PS Solomon. Just the surface: advances in the discovery and characterization of necrotrophic wheat effectors. Curr. Opin. Microbiol. 2018 46:14–8. 10.1016/J.MIB.2018.01.019.

19. Friesen TL, EH Stukenbrock, Z Liu, et al. Emergence ofa new disease asa result of interspecific virulence gene transfer. Nat. Genet. 2006 388 2006 38:953–6. 10.1038/ng1839.

20. Vleeshouwers VGAA, RP Oliver. Effectors as tools in disease resistance breeding against biotrophic, hemibiotrophic, and necrotrophic plant pathogens. Mol. Plant. Microbe Interact. 2014 27:196–206. 10.1094/MPMI-10-13-0313-IA.

21. Francki MG, E Walker, CJ McMullan, et al. Multi-Location Evaluation of Global Wheat Lines Reveal Multiple QTL for Adult Plant Resistance to Septoria Nodorum Blotch (SNB) Detected in Specific Environments and in Response to Different Isolates. Front. Plant Sci. 2020 11. 10.3389/fpls.2020.00771.

22. Shankar M, D Mather, H Golzar, et al. Stacking yellow spot resistance genes into fixed lines results in genetic gain. 2015. https://www.researchgate.net/publication/305748820.

23. Peters Haugrud AR, G Shi, S Seneviratne, et al. Genome-wide association mapping of resistance to the foliar diseases septoria nodorum blotch and tan spot in a global winter wheat collection. Mol. Breed. New Strateg. Plant Improv. 2023 43:54. 10.1007/s11032-023-01400-5.

24. Beard C, G Thomas, A Hills, et al. Yellow spot and septoria nodorum blotch and their management in wheat. 2024. https://library.dpird.wa.gov.au/fc_factsheets/70.

25. Friskop A, Z Liu. Fungal Leaf Spot Diseases of Wheat: Tan spot, Septoria/Stagonospora nodorum blotch and Septoria tritici blotch (PP1249). 2021. www.ag.ndsu.edu/extension.

26. Long M, M Hartley, RJ Morris, et al. Classification of wheat diseases using deep learning networks with field and glasshouse images. Plant Pathol. 2023 72:536–47. 10.1111/ppa.13684.

27. Abdullah AS, C Turo, CS Moffat, et al. Real-time PCR for diagnosing and quantifying co-infection by two globally distributed fungal pathogens of wheat. Front. Plant Sci. 2018 9:1086–1086. 10.3389/FPLS.2018.01086/BIBTEX.

28. Fu H, J Jiang, MW Harding, et al. Development of a triplex qPCR system for the detection of Parastagonospora nodorum, Zymoseptoria tritici and Pyrenophora tritici-repentis from wheat. Plant Dis. 2025. https://apsjournals.apsnet.org/doi/10.1094/PDIS-01-25-0003-SR (18 August 2025, date last accessed).

29. Moffat CS, PT See, RP Oliver. Leaf yellowing of the wheat cultivar Mace in the absence of yellow spot disease. Australas. Plant Pathol. 2015 44:161–6. 10.1007/s13313-014-0335-2.

30. Hall AT, AM Zovanyi, DR Christensen, et al. Evaluation of Inhibitor-Resistant Real-Time PCR Methods for Diagnostics in Clinical and Environmental Samples. PLOS ONE 2013 8:e73845. 10.1371/journal.pone.0073845.

31. Huggett JF, CA Foy, V Benes, et al. The digital MIQE guidelines: Minimum information for publication of quantitative digital PCR experiments. Clin. Chem. 2013 59:892–902. 10.1373/clinchem.2013.206375.

32. Solomon PS, SW Thomas, P Spanu, et al. The utilisation of di/tripeptides by Stagonospora nodorum is dispensable for wheat infection. Physiol. Mol. Plant Pathol. 2003 63:191–9. 10.1016/j.pmpp.2003.12.003.

33. Moolhuijzen P, PT See, CS Moffat. A new PacBio genome sequence of an Australian Pyrenophora tritici-repentis race 1 isolate. BMCRes. Notes 2019 12:642–642. 10.1186/s13104-019-4681-6.

34. Rybak K, PT See, HTT Phan, et al. Afunctionally conserved Zn2Cys6 binuclear cluster transcription factor class regulates necrotrophic effector gene expression and host-specific virulence of two major Pleosporales fungal pathogens of wheat. Mol. Plant Pathol. 2017 18:420–34. 10.1111/mpp.12511.

35. Liu ZH, JD Faris, SW Meinhardt, et al. Genetic and physical mapping of a gene conditioning sensitivity in wheat to apartially purified host-selective toxin produced by Stagonospora nodorum. Phytopathology 2004 94:1056–60.

36. Jacques S, L Lenzo, K Stevens, et al. An optimized sporulation method for the wheat fungal pathogen Pyrenophora tritici-repentis. Plant Methods 2021 17:52–52. 10.1186/s13007-021-00751-4.

37. Abràmoff MD, PJ Magalhães, SJ Ram. Image processing with ImageJ. Biophotonics Int. 2004 11:36–42.

38. Jones DAB, K Rybak, M Hossain, et al. Repeat-induced point mutations driving Parastagonospora nodorum genomic diversity are balanced by selection against non-synonymous mutations. *Commun*. Biol. 2024 7:1614. 10.1038/s42003-024-07327-7.

39. Syme RA, K-C Tan, K Rybak, et al. Pan-Parastagonospora comparative genome analysis—effector prediction and genome evolution. Genome Biol. Evol. 2018 10:2443–57.

40. Moolhuijzen PM, PT See, G Shi, et al. A global pangenome for the wheat fungal pathogen Pyrenophora tritici-repentis and prediction of effector protein structural homology. *Microb*. Genomics 2022 8:mgen000872. 10.1099/mgen.0.000872.

41. R Core Team. R: A Language and Environment for Statistical Computing. 2025.

42. Keene ON. The log transformation is special. Stat. Med. 1995 14:811–9. 10.1002/sim.4780140810.

43. Suffert F, G Delestre, F Carpentier, et al. Fashionably late partners have more fruitful encounters: Impact of the timing of co-infection and pathogenicity on sexual reproduction in Zymoseptoria tritici. Fungal Genet. Biol. 2016 92:40–9. 10.1016/J.FGB.2016.05.004.

44. Barrett LG, M Zala, A Mikaberidze, et al. Mixed infections alter transmission potential in a fungal plant pathogen. Environ. Microbiol. 2021 23:2315–30. 10.1111/1462-2920.15417.

45. John E, S Jacques, HTT Phan, et al. Variability in an effector gene promoter of a necrotrophic fungal pathogen dictatesepistasisand effector-triggered susceptibility inwheat. PLOSPathog.2022 18:e1010149.

46. Bathgate JA, R Loughman. Ascosporesare asource of inoculum of Phaeosphaeria nodorum,P. Avenaria f. sp. Avenaria and Mycosphaerella graminicola in Western Australia. Australas. Plant Pathol. 2001 30:317–22. 10.1071/AP01043.

47. Galloway J, M Salam, P Payne. Less yellow spot in wheat: progress towardsa decision support tool that will predict whenyellow spot sporesarereleased from the previous season stubble. 2016.

48. Benslimane H, S Aouali, A Khalfi, et al. In vitro morphological characteristics of Pyrenophora tritici-repentis isolates from several Algerian agro-ecological zones. Plant Pathol. J. 2017 33:109–17. 10.5423/PPJ.OA.09.2015.0189.

49. Muria-Gonzalez MJ, Y Yeng, S Breen, et al. Volatile Molecules Secreted by the Wheat Pathogen Parastagonospora nodorum Are Involved in Development and Phytotoxicity. Front. Microbiol. 2020 11. 10.3389/fmicb.2020.00466.

50. Seybold H, TJ Demetrowitsch, MA Hassani, et al. A fungal pathogen induces systemic susceptibility and systemic shifts in wheat metabolome and microbiome composition. Nat. Commun. 2020 11. https://pubmed.ncbi.nlm.nih.gov/32313046/.

51. Balint-Kurti P. The plant hypersensitive response: concepts, control and consequences. Mol. Plant Pathol. 2019 20:1163–78. 10.1111/MPP.12821.

52. Glazebrook J. Contrasting mechanisms of defense against biotrophic and necrotrophic pathogens. Annu. Rev. Phytopathol. 2005 43:205–27. 10.1146/annurev.phyto.43.040204.135923.

53. Tan KC, RP Oliver. Regulation of proteinaceous effector expression in phytopathogenic fungi. PLoS Pathog. 2017 13. https://pubmed.ncbi.nlm.nih.gov/28426760/.

54. Kariyawasam GK, JK Richards, NA Wyatt, et al. The Parastagonospora nodorum necrotrophic effector SnTox5 targetsthe wheatgene Snn5 and facilitatesentry into the leafmesophyll. New Phytol.2022 233:409–26. 10.1111/nph.17602.

55. Richards JK, GK Kariyawasam, S Seneviratne, et al. A triple threat: the Parastagonospora nodorum SnTox267 effector exploits three distinct host genetic factors to cause disease in wheat. New Phytol. 2022 233:427–42. 10.1111/nph.17601.

56. Snelders NC, GC Petti, GCM van den Berg, et al. An ancient antimicrobial protein co-opted by a fungal plant pathogen for in planta mycobiome manipulation. Proc. Natl. Acad. Sci. 2021 118:e2110968118– e2110968118. 10.1073/pnas.2110968118.

57. Liu Z, Y Gao, YM Kim, et al. SnTox1, a Parastagonospora nodorum necrotrophic effector, is a dual-function protein that facilitates infection while protecting from wheat-produced chitinases. New Phytol. 2016 211:1052–64. 10.1111/nph.13959.

58. Chooi YH, MJ Muria-Gonzalez, PS Solomon. A genome-wide survey of the secondary metabolite biosynthesis genes in the wheat pathogen Parastagonospora nodorum. Mycology 2014 5:192–206. 10.1080/21501203.2014.928386.

59. Chooi YH, C Krill, RA Barrow, et al. An In planta-expressed polyketide synthase produces (R)-mellein in the wheat pathogen Parastagonospora nodorum. Appl. Environ. Microbiol. 2015 81:177–86. 10.1128/AEM.02745-14.

60. Chooi YH, G Zhang, J Hu, et al. Functional genomics-guided discovery of a light-activated phytotoxin in the wheat pathogen Parastagonospora nodorum via pathway activation. Environ. Microbiol. 2017 19:1975–86. 10.1111/1462-2920.13711.

61. Robey MT, LK Caesar, MT Drott, et al. Aninterpreted atlasofbiosynthetic geneclusters from 1,000 fungal genomes. Proc. Natl. Acad. Sci. 2021 118:e2020230118. 10.1073/pnas.2020230118.

62. Bock CH, JGA Barbedo, EM Del Ponte, et al. From visual estimates to fully automated sensor-based measurements of plant disease severity: status and challenges for improving accuracy. Phytopathol. Res. 2020 2. 10.1186/s42483-020-00049-8.

63. Faris JD, Z Zhang, H Lu, et al. A unique wheat disease resistance-like gene governs effector-triggered susceptibility to necrotrophic pathogens. Proc. Natl. Acad. Sci. 2010 107:13544–9. 10.1073/pnas.1004090107.

64. Shackley B, C Zaicou-Kunesch, H Dhammu, et al. Wheat Variety Guide for WA Wheat variety guide for WA 2014. 2014.

65. Shackley B, D Nicol, J Curry, et al. Department of Primary Industries and Regional Development Crop Sowing Guide 2022. 2022.

66. Bertazzoni S, DAB Jones, HT Phan, et al. Chromosome-level genome assembly and manually-curated proteome of model necrotroph Parastagonospora nodorum Sn15 reveals a genome-wide trove of candidate effector homologs, and redundancy of virulence-related functions within an accessory chromosome. BMC Genomics 2021 22:382–382. 10.1186/s12864-021-07699-8.

67. McDonald MC, AP Taranto, E Hill, et al. Transposon-mediated horizontal transfer of the host-specific virulence protein ToxA between three fungal wheat pathogens. mBio 2019 10. 10.1128/mBio.01515-19.

