## Supplementary Figures for "Fair-weather friends: Unequal partnerships between *Parastagonospora nodorum* and *Pyrenophora tritici-repentis* define disease dynamics in wheat"

### Supplementary Tables

***Table S1: HSV values used to generate coverage masks for infected tissue.***

| Tissue Type | H_min | H_max | S_min | S_max | V_min | V_max |
|---|---|---|---|---|---|---|
| Background | 90 | 255 | 0 | 255 | 0 | 255 |
| Healthy | 43 | 90 | 0 | 255 | 0 | 255 |
| Chlorotic | 30 | 43 | 0 | 255 | 0 | 255 |
| Necrotic | 0 | 30 | 0 | 255 | 0 | 255 |

***Table S2 Strains used in this study***

| Species | Strain | Reference |
|---|---|---|
| *Parastagonospora nodorum* | SN15 | [66] |
| *Pyrenophora tritici-repentis* | M4 | [33] |
| *Bipolaris sorokiniana* | CS10 | [67] |
| *Fusarium graminearum* | WAC11354 | *Unpublished* |
| *Pyrenophora teres teres* | W1-1 | [39] |
| *Pyrenophora teres maculata* | SG-1 | [39] |
| *Alternaria alternata* | | *Unpublished* |

### Supplementary Figures

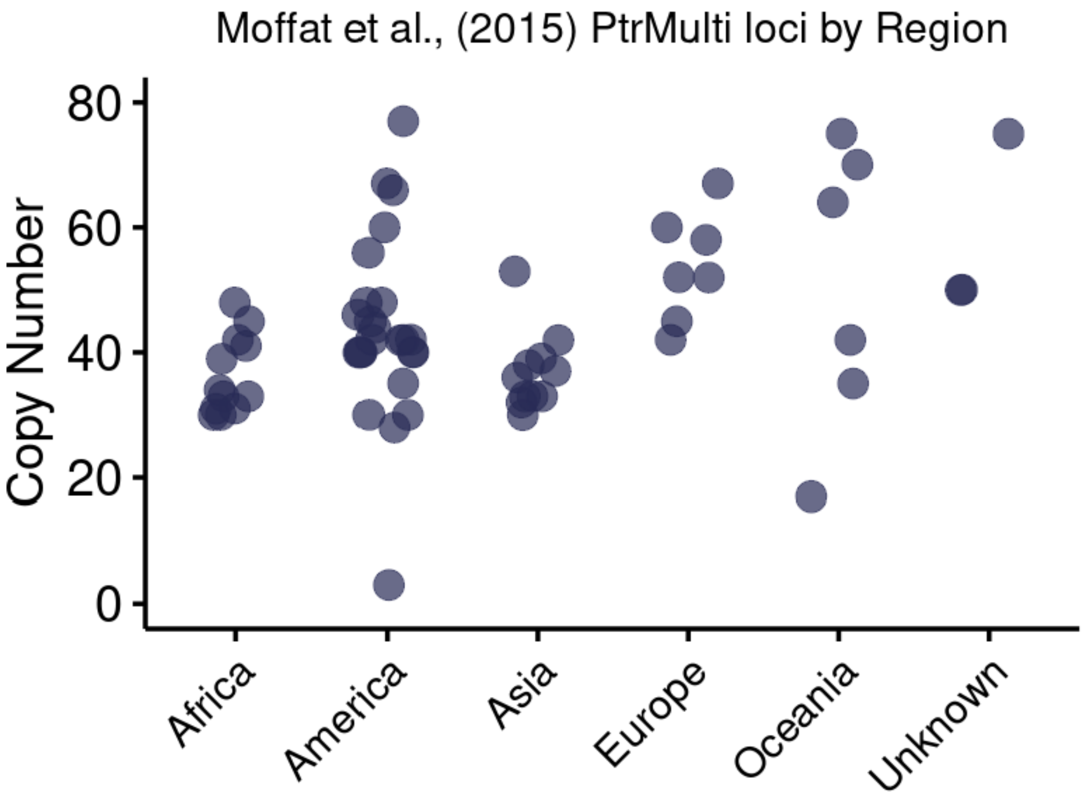
Moffat et al., (2015) PtrMulti loci by Region
Copy Number
0
20
40
60
80
Africa
America
Asia
Europe
Oceania
Unknown

***Figure S1. Copy number variation***

Copy numbers determined by BLASTN query of the PtrMulti locus for perfect matches (100% identity) against publicly available genome assemblies of *P. tritici-repentis*. Each point represents one genome assembly. Copy numbers indicate the total number of perfect sequence matches identified within each assembly.

A

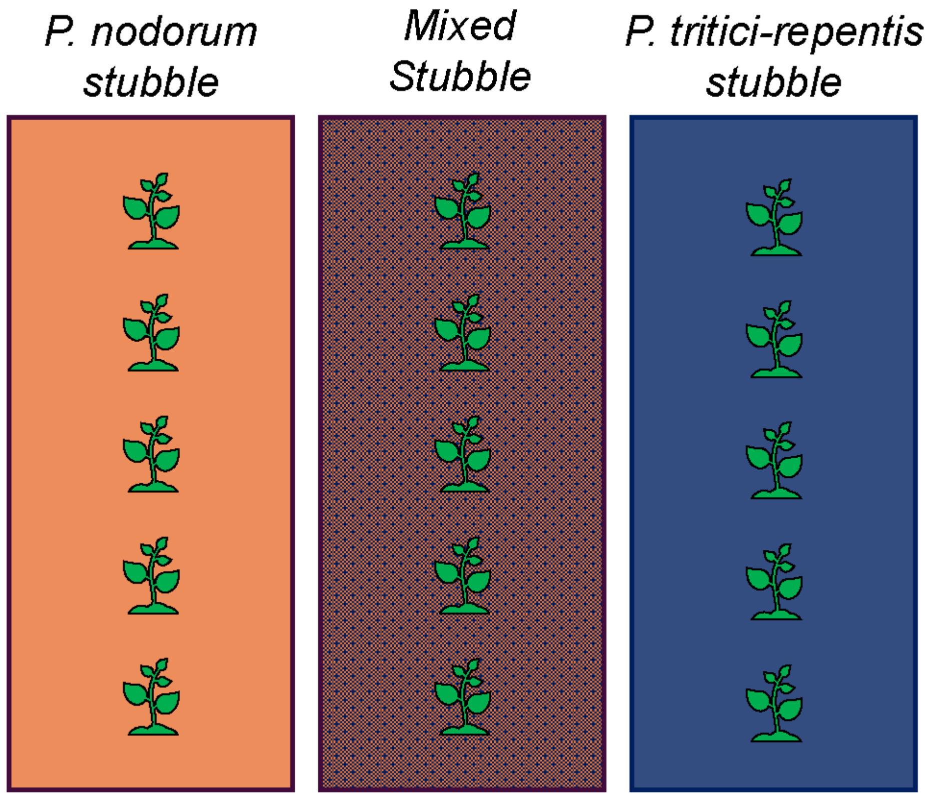

*P. nodorum* stubble
*Mixed* Stubble
*P. tritici-repentis* stubble

***Figure S2: Field trial design***

Shows field trial design for controlled inoculum studies. Single inoculum plots were placed on the edges of the field, while the mixed inoculum was placed in the middle.

A

$$DNA_{(ug)} = \beta \times Tissue_{(mg)}$$

$$Copies_{(aT)} = \alpha \times DNA_{(ng)}$$

$$\therefore$$

$$Tissue_{(ug)} = \frac{Copies_{(aT)}}{\alpha\beta}$$

B

$$Tissue_{(ug)} = \frac{Copies_{(aT)}}{0.16 \times 4825}$$

$$Tissue_{(ug)} = 1.29 \times Copies_{(aT)}$$

C

$$Tissue_{(ug)} = \frac{Copies_{(aT)}}{0.059 \times 4974}$$

$$Tissue_{(ug)} = 3.36 \times Copies_{(aT)}$$

***Figure S3. Integrated biomass model***

Linear Equations for calculating biomass, given ddPCR target copies. **(A)** shows the general form, where α = the slope of DNA(μg) over Tissue(mg), and β = the slope of ddPCR copies over DNA(ng), while **(B)** and **(C)** show specific forms for *P. tritici-repentis* (blue), and *P. nodorum* (orange).

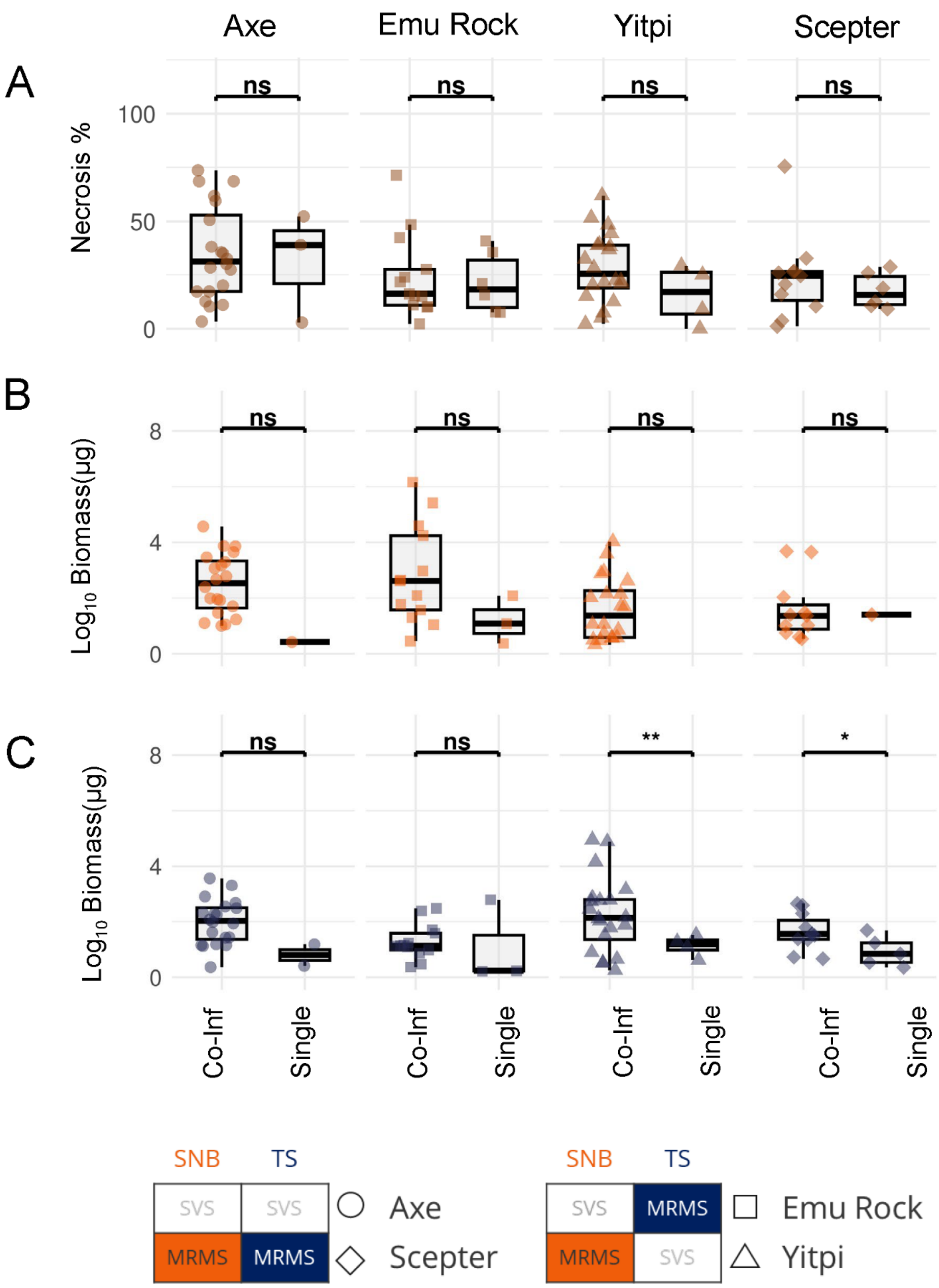
Axe
Emu Rock
Yitpi
Scepter
A
Necrosis %
100
50
0
ns
B
Log10 Biomass(µg)
8
4
0
C
**
*
Co-Inf
Single
SNB
TS
SVS
MRMS
Axe
Scepter
Emu Rock
Yitpi

***Figure S4: P. tritici-repentis infected stubble***

**(A)** disease severity (necrosis %, brown), **(B)** *P. nodorum* biomass (orange), and **(C)** *P. tritici-repentis* (blue) biomass across four wheat lines: Axe (SVS to SNB and TS), Emu Rock (SVS to SNB, MRMS to TS), Scepter (MRMS to SNB and TS), and Yitpi (MRMS to SNB SVS to TS). Points represent individual wheat leaves with shapes indicating cultivar (circles = Axe, squares = Emu, diamonds = Scepter, triangles = Yitpi). Statistical significance of differences between co-infection and single infection within each wheat line indicated above brackets (ns = not significant, * $p < 0.05$, ** $p < 0.01$, *** $p < 0.001$).

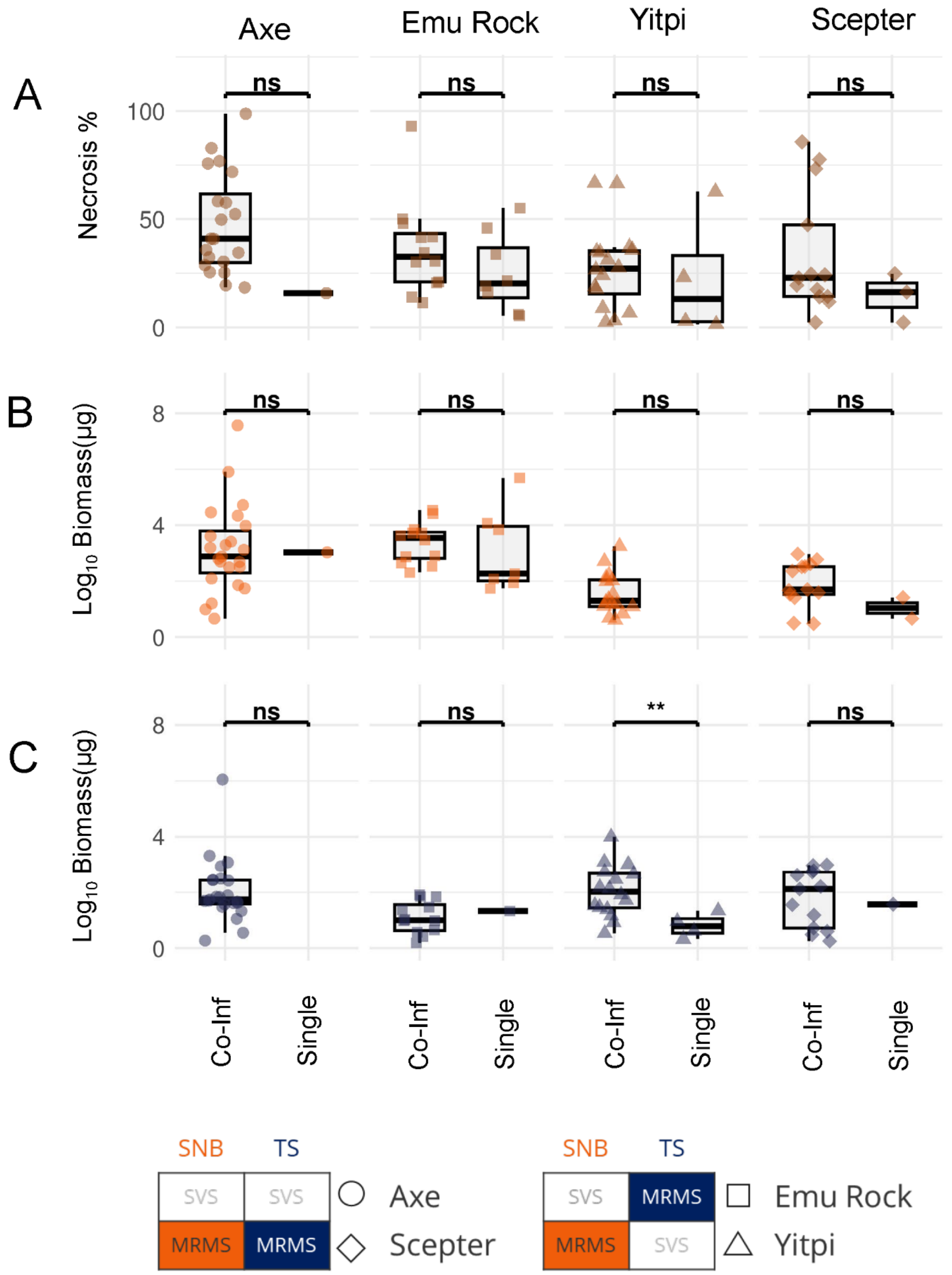

Axe
Emu Rock
Yitpi
Scepter
A
Necrosis %
100
50
0
ns
B
Log10 Biomass(μg)
8
4
0
C
**
Co-Inf
Single
SNB
TS
SVS
MRMS
Axe
Scepter
Emu Rock
Yitpi

***Figure S5: Mixed P. nodorum and P. tritici-repentis infected stubble***

**(A)** disease severity (necrosis %, brown), **(B)** *P. nodorum* biomass (orange), and **(C)** *P. tritici-repentis* (blue) biomass across four wheat lines: Axe (SVS to SNB and TS), Emu Rock (SVS to SNB, MRMS to TS), Scepter (MRMS to SNB and TS), and Yitpi (MRMS to SNB SVS to TS). Points represent individual wheat leaves with shapes indicating cultivar (circles = Axe, squares = Emu, diamonds = Scepter, triangles = Yitpi). Statistical significance of differences between co-infection and single infection within each wheat line indicated above brackets (ns = not significant, * $p < 0.05$, ** $p < 0.01$, *** $p < 0.001$).

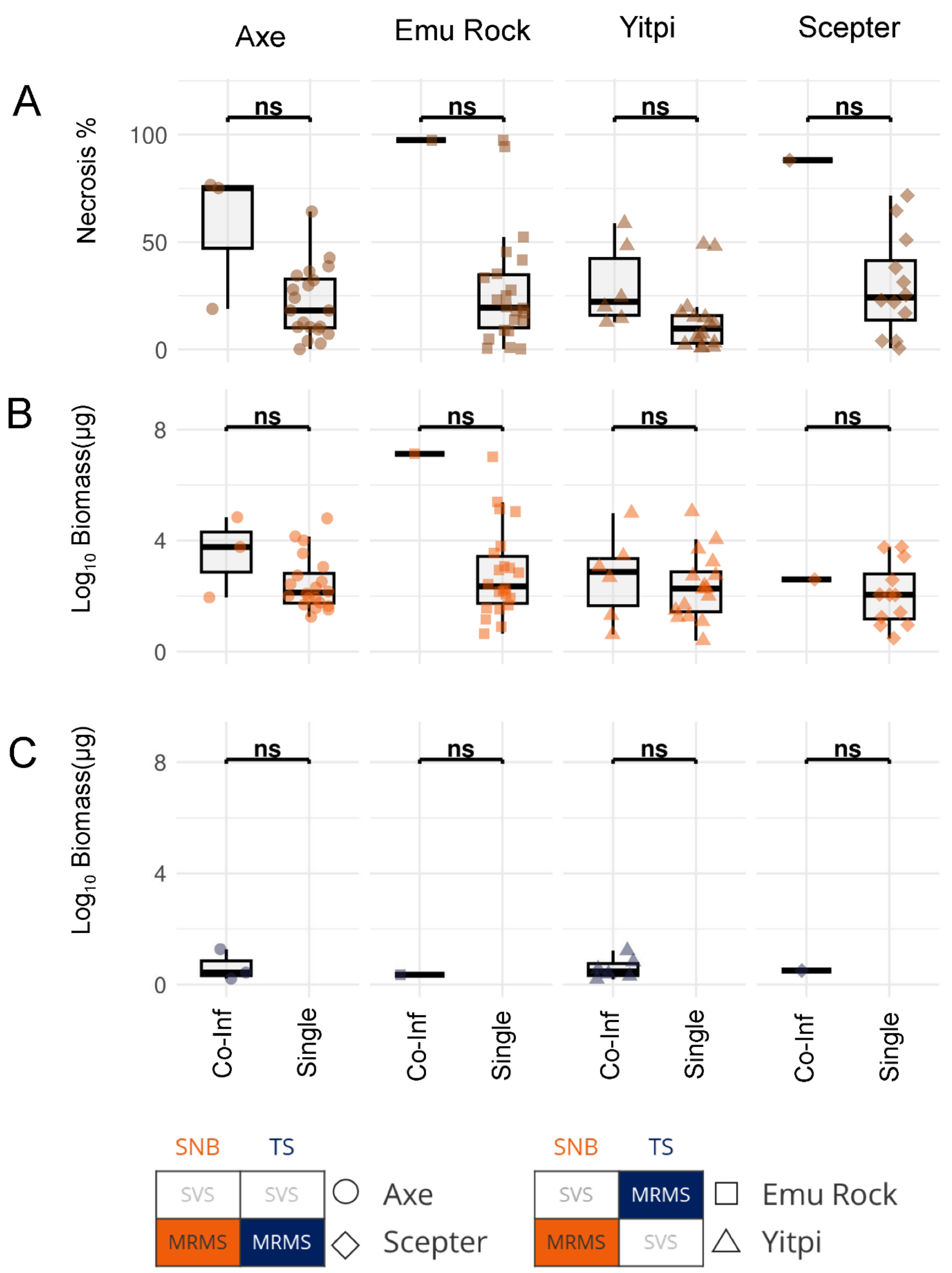
Axe
Emu Rock
Yitpi
Scepter
A
Necrosis %
100
50
0
ns
B
Log10 Biomass(µg)
8
4
0
C
Co-Inf
Single
SNB
TS
SVS
MRMS
Axe
Scepter
Emu Rock
Yitpi

***Figure S6: P. nodorum infected stubble***

**(A)** disease severity (necrosis %, brown), **(B)** *P. nodorum* biomass (orange), and **(C)** *P. tritici-repentis* (blue) biomass across four wheat lines: Axe (SVS to SNB and TS), Emu Rock (SVS to SNB, MRMS to TS), Scepter (MRMS to SNB and TS), and Yitpi (MRMS to SNB SVS to TS). Points represent individual wheat leaves with shapes indicating cultivar (circles = Axe, squares = Emu, diamonds = Scepter, triangles = Yitpi). Statistical significance of differences between co-infection and single infection within each wheat line indicated above brackets (ns = not significant, * $p < 0.05$, ** $p < 0.01$, *** $p < 0.001$).
